# Xenium In Situ Profiling Uncovers HSPG-Dependent SULF1/VEGFR2 Signaling Mediating Vascular Remodeling in Moyamoya Disease

**DOI:** 10.64898/2026.04.28.721514

**Authors:** Yaoren Chang, Xiaofan Yu, Talha Ahmed, Yuanli Zhao, Shihao He, Xun Ye

**Affiliations:** Department of Neurosurgery, Beijing Tiantan Hospital, Capital Medical University, Beijing, 100070, China; China National Clinical Research Center for Neurological Diseases, Beijing, 100070, China; Department of Neurosurgery, Beijing Chao-Yang Hospital, Capital Medical University; Beijing, 100020, China; Department of Pathology, Johns Hopkins School of Medicine, Baltimore, Maryland, 21205, USA; Department of Neurosurgery, Peking Union Medical College Hospital, Peking Union Medical College and Chinese Academy of Medical Sciences; Beijing, 100730, China

**Keywords:** Moyamoya disease, Xenium, SULF1, Cell migration, Intimal thickening, spatial transcriptomics, angiogenesis

## Abstract

**Background:** Moyamoya disease (MMD) is characterized by progressive arterial stenosis and abnormal collateral formation, but the spatial organization of vessel-wall abnormalities remains incompletely understood.

**Methods:** We combined Xenium in situ spatial transcriptomics and multiplex immunofluorescence in superficial temporal artery samples from patients with MMD and controls, and performed gain- and loss-of-function experiments in human brain microvascular endothelial cells (HBMECs). Western blotting, quantitative real-time polymerase chain reaction (qRT-PCR), tube-formation, Transwell migration, and cell scratch assays were used to assess signaling and endothelial phenotypes.

**Results:** MMD vascular tissue showed intimal hyperplasia, altered spatial cellular architecture, and enrichment of extracellular matrix- and proteoglycan-related programs, with upregulation of sulfatase 1 (SULF1). In HBMECs, SULF1 knockdown reduced, whereas SULF1 overexpression enhanced, vascular endothelial growth factor A165 (VEGF-A165)-induced vascular endothelial growth factor receptor 2 (VEGFR2), extracellular signal-regulated kinase 1/2 (ERK1/2), and protein kinase B (AKT) phosphorylation, migration, tube formation, and angiogenesis- and adhesion-related gene expression. Heparinase III attenuated the signaling effects associated with SULF1 overexpression.

**Conclusion:** These findings suggest that SULF1-associated extracellular matrix alterations may contribute to local vessel-wall remodeling and enhanced endothelial responsiveness in MMD.

**Graphical Abstract:** 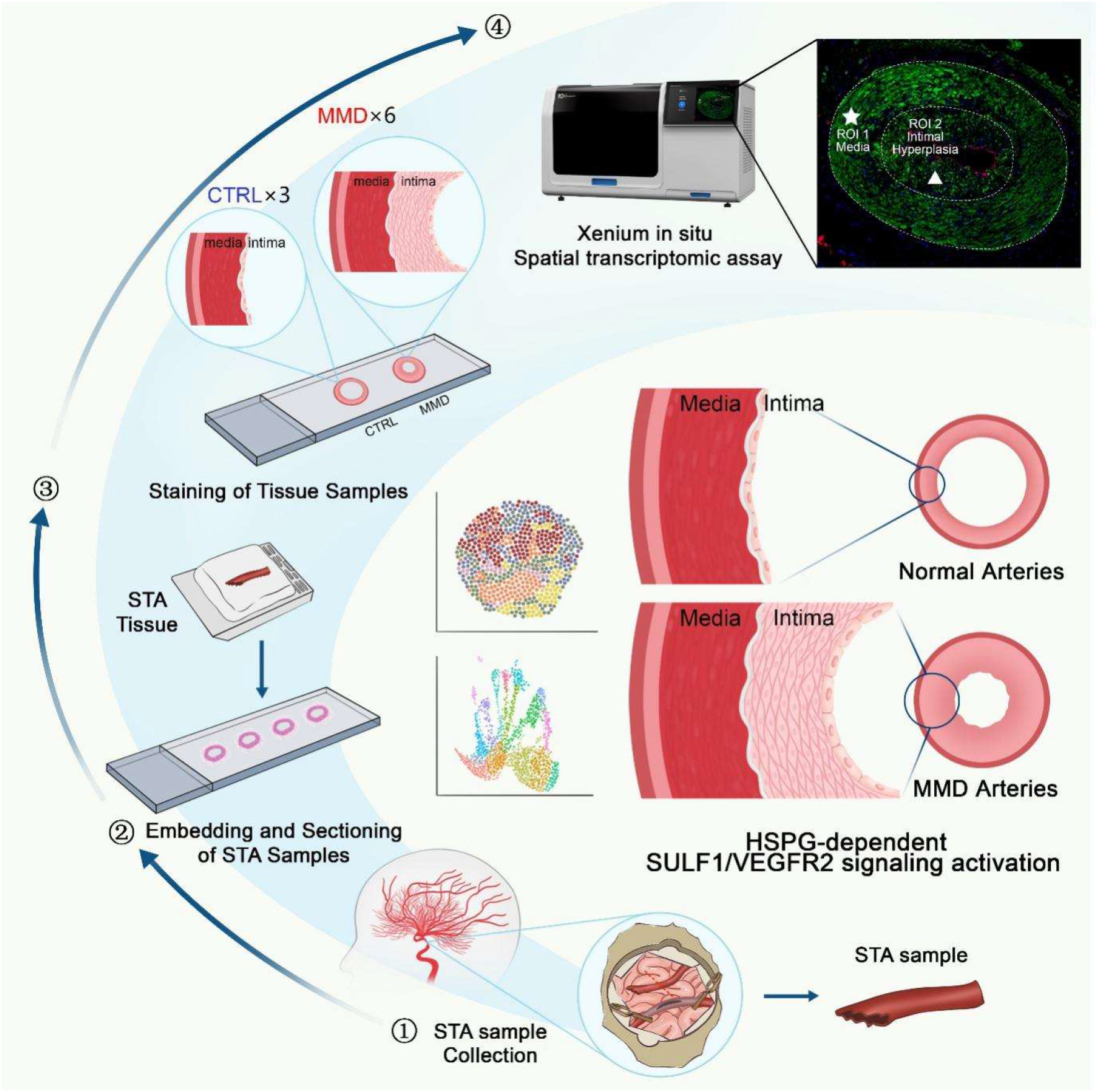

## Introduction

Moyamoya disease (MMD) is a chronic occlusive cerebrovascular disorder characterized by progressive stenosis or occlusion of the terminal internal carotid artery and its proximal branches, together with the formation of an abnormal collateral vascular network at the base of the brain ^1–3^. MMD may present with transient ischemic attacks, cerebral infarction, intracranial hemorrhage, cognitive impairment, and progressive neurological dysfunction in both pediatric and adult patients ^1,2^. The affected vessels typically show fibrocellular intimal thickening, irregular or duplicated internal elastic lamina, medial thinning, and abnormal vascular remodeling, indicating that MMD is not simply a disorder of luminal narrowing but a dynamic vasculopathy involving complex changes in the vessel wall ^1,2,4^. Despite growing advances in genetics and studies of vessel-wall remodeling, the mechanisms driving lesion formation and progression in MMD remain incompletely understood, and surgical revascularization is still the mainstay of treatment ^2,4^.

Previous genetic, epigenetic, and molecular studies have linked MMD to multiple abnormalities related to vascular remodeling and angiogenesis. Genome-wide association analyses linked *ring finger protein 213 (RNF213)*, particularly the p.R4810K variant, to disease susceptibility in East Asian populations, and additional loci such as *actin alpha 2 (ACTA2), diaphanous-related formin 1 (DIAPH1)*, and *HLA*-related variants have also been implicated in vascular abnormalities ^4,5^. Sung et al. used an Illumina 450K methylation array in endothelial cells and found that sortilin 1 (SORT1) promoter hypomethylation was associated with enhanced angiogenic activity ^6^. An epigenome-wide association study of adults with ischemic MMD used an 850K methylation array in whole blood and identified differentially methylated loci enriched in pathways related to angiogenesis and cell growth ^7^. Ye et al. combined *Rnf213*-deficient mice and *rnf213*-deficient zebrafish, *ring finger protein 213 (RNF213)* knockdown in human brain microvascular endothelial cells (HBMECs), and transcriptomic analyses, and found that *ring finger protein 213 (RNF213)* loss of function promoted endothelial proliferation, migration, and tube formation with activation of Hippo-Yes-associated protein/transcriptional coactivator with PDZ-binding motif (YAP/TAZ) signaling ^8^. He et al. analyzed circulating small extracellular vesicle (sEV)/exosomal RNAs and identified dysregulated circular RNAs (circRNAs) associated with angiogenesis and vascular occlusion, whereas a data-independent acquisition (DIA)-based serum proteomic study further linked filamin A (FLNA) and zyxin (ZYX) upregulation to abnormal angiogenesis and cytoskeletal changes in MMD ^9,10^. More recently, a single-cell transcriptomic study of superficial temporal artery tissue further identified vascular and immune cell heterogeneity in MMD, and suggested an increased proportion of smooth muscle cells together with abnormal interactions between immune cells and vascular cells ^11^. These findings have outlined molecular and cellular abnormalities in MMD, but they still provide limited direct information on how such changes are organized within the diseased vessel wall in situ.

Spatial transcriptomic methods can retain tissue context and align molecular information with histological architecture, making them more suitable for resolving lesion-associated microenvironments ^12,13^. Fan et al. integrated single-cell RNA sequencing with spatial transcriptomic data to analyze tertiary lymphoid structure-associated regions in breast cancer and identified 18 major cell types, macrophage lineage transitions, and fibroblast-associated C-X-C motif chemokine ligand 12 (CXCL12)-C-X-C chemokine receptor type 4 (CXCR4) signaling related to lymphocyte recruitment ^14^. Campos et al. used GeoMx and CosMx on serial sections of atherosclerotic coronary arteries to distinguish immune-cell distribution and neighborhood relationships across different layers of the vessel wall ^15^. Yet the choice of platform remains question-dependent. Sequencing-based platforms are useful for broader discovery and larger-area tissue mapping, but in structurally compact vessel-wall lesions their effective resolution may still rely on spot size, binning, or deconvolution ^12,13^. Salas et al. compared Xenium with other spatial transcriptomic technologies and showed that Xenium can detect targeted transcripts in situ at subcellular resolution, making it more suitable for resolving local vessel-wall architecture and closely apposed cell populations in MMD ^16^.

In this study, we applied Xenium in situ spatial transcriptomics to MMD vascular tissue to characterize the spatial cellular architecture and lesion-associated microenvironment. We further integrated histological assessment and multiplex immunofluorescence to validate spatial cellular distribution in tissue sections, and used Western blotting, quantitative real-time polymerase chain reaction (qRT-PCR), cell scratch assays, and tube-formation assays to examine relevant molecular and functional features. We aimed to define the spatial organization and local distribution of disease-associated changes within the MMD vessel wall.

## Methods

### Participants and tissue collection

Patients with MMD were clinically diagnosed on the basis of medical history and neuroradiological examinations according to the diagnostic guidelines. Neuroradiological evaluations included cerebral digital subtraction angiography, magnetic resonance imaging, and functional regional cerebral blood flow studies. Clinical and imaging data were obtained from medical records and the Picture Archiving and Communication System, respectively. Superficial temporal artery (STA) segments measuring 1–3 mm were collected from patients with MMD during revascularization surgery, including direct or combined bypass procedures. Control STA samples were obtained from patients undergoing craniotomy for epilepsy management. Written informed consent was obtained from all participants or their legal guardians. This study was approved by the Institutional Ethics Committee Of Beijing Tiantan Hospital (KYSQ2024-248-01).

### Multiplex immunofluorescence staining and morphometric analysis

Vascular sections were subjected to multiplex immunofluorescence staining using alpha-smooth muscle actin (αSMA) and cluster of differentiation 31 (CD31) to evaluate vessel-wall structure and endothelial distribution. All images were acquired under identical imaging settings. The intimal and medial areas were measured separately, and the intima-to-media area ratio was calculated for group comparison.

### Xenium in situ spatial transcriptomic assay

Fresh-frozen (FF) or formalin-fixed paraffin-embedded (FFPE) vascular tissue sections were mounted within the effective capture area of Xenium slides, which measured 10.45 mm × 22.45 mm. Before assay initiation, the sections underwent 4′,6-diamidino-2-phenylindole (DAPI) staining, hematoxylin and eosin (H&E) staining, and RNA quality assessment to evaluate nuclear quality, tissue morphology, and RNA integrity, and regions of interest were selected accordingly. FF sections were subjected to fixation and permeabilization, whereas FFPE sections underwent deparaffinization, rehydration, and decrosslinking to expose RNA molecules for probe hybridization. Circularizable DNA probes targeting the selected gene panel were hybridized to the tissue overnight in a thermal cycler for 16–24 h, followed by probe ligation and in situ rolling-circle amplification to generate gene-specific spatial barcode sequences. After nuclear restaining with DAPI, the slides were loaded onto the Xenium Analyzer (10x Genomics, Pleasanton, CA, USA) for repeated cycles of fluorescent probe hybridization, imaging, and probe removal to decode transcript identities and spatial locations. When applicable, multimodal cell segmentation was performed using membrane and cytoplasmic protein antibodies together with cytoplasmic 18S RNA probes.

### Xenium data processing, clustering, and cell-type annotation

Raw Xenium data were processed using Xenium Ranger (version 3.1.0.4; 10x Genomics, Pleasanton, CA, USA) with default parameters. The Xenium-derived expression matrices and cell-boundary files were imported into Seurat V5 for downstream analysis. Cells with fewer than 5 detected genes and genes expressed in fewer than 5 cells were excluded. Additional quality-control thresholds were defined according to the distributions of nFeature_RNA, nCount_RNA, and log10GenesPerUMI to remove low-quality cells. Because of the large scale of the dataset, BPCells was used to reduce computational burden. A sketch-based workflow was then applied, and 50,000 cells were subsampled using the LeverageScore method for initial clustering. Data normalization was performed using SCTransform, followed by identification of highly variable genes, data scaling, and principal component analysis. The first 15 principal components were used for downstream analysis. Shared nearest-neighbor graphs were generated with FindNeighbors, and clusters were identified with FindClusters using a resolution of 1.8. Uniform manifold approximation and projection (UMAP) was used for visualization. Cluster labels derived from the sketched dataset were projected back to the full dataset using ProjectData. Cell types were annotated according to canonical marker-gene expression patterns, aided by DotPlot and Heatmap visualization. To refine subcluster annotation, major cell-type subsets were separated and reanalyzed independently without sketch-based subsampling, using the full subset for normalization, dimensionality reduction, reclustering, and marker-based annotation.

### Differential expression and functional enrichment analyses

Differential expression analyses were performed for overall group comparisons, cell-type-specific comparisons, and selected subgroup or spatial-region comparisons. Differentially expressed genes were ranked according to adjusted P values, average log2 fold changes, and the proportions of expressing cells. Genes from relevant comparisons were subsequently subjected to Kyoto Encyclopedia of Genes and Genomes and Gene Ontology enrichment analyses to identify lesion-associated biological processes and molecular functions.

### Cellular neighborhood analysis

Cellular neighborhood analysis was performed using cell-type labels, sample-group information, and spatial coordinates as input. The nn2 function from the RANN package was used to identify the 20 nearest neighbors for each cell. Cells were then grouped into 20 cellular neighborhoods using the kmeans function on the basis of neighborhood composition. The composition and spatial distribution of each cellular neighborhood were compared between groups.

### Cell culture

HBMECs were purchased from ScienCell (Carlsbad, CA, USA) and maintained in endothelial cell medium (cat. no. 1001; Shanghai Zhong Qiao Xin Zhou Biotechnology, Shanghai, China) supplemented with 5% fetal bovine serum (FBS; SH30070.03; Hyclone, Logan, UT, USA), 1% endothelial cell growth supplement (cat. no. 1052; Shanghai Zhong Qiao Xin Zhou Biotechnology, Shanghai, China), and 1% penicillin-streptomycin (15140148; KeyGEN, Nanjing, China). HBMECs were dissociated using trypsin-EDTA solution (KGY0012; KeyGEN, Nanjing, China). HBMECs were cultured at 37°C in a humidified incubator with 5% CO2 and passaged at approximately 90% confluence.

### Lentiviral transfection and experimental grouping

For knockdown experiments, HBMECs were transfected with the negative-control lentivirus CON313 or LV-SULF1-RNAi targeting sulfatase 1 (SULF1). After validation, LV-SULF1-RNAi (PSC60698-1) was selected for subsequent experiments. For overexpression experiments, HBMECs were transfected with the control lentivirus CON335 or LV-SULF1 (KL25472-1). Lentiviral infection was performed with HitransG P and HitransG A viral transduction reagents provided by the client. For signaling assays, HBMECs were cultured overnight in medium containing 1% fetal bovine serum (FBS) and then stimulated with vascular endothelial growth factor A165 (VEGF-A165) protein (HY-P70458A; MedChemExpress [MCE], New Jersey, USA) for 0, 10, and 30 min. For heparinase III pretreatment experiments, HBMECs were cultured in medium containing 1% FBS for 24 h, pretreated with heparinase III (HY-P2953; MCE, New Jersey, USA) at 20 mU for 2 h, and then stimulated with VEGF-A165 (50 ng/mL) for 0, 10, and 30 min. For functional assays and qRT-PCR, transfected HBMECs were treated with VEGF-A165 at a final concentration of 10 ng/mL. Unless otherwise specified, all cell experiments were performed with three independent replicates (N = 3).

### Tube-formation assay

Matrigel (M8370; Solarbio Science & Technology, Beijing, China) was thawed at 4°C overnight and added to 96-well plates at 50 μL per well, followed by incubation at 37°C for 30 min to allow solidification. Treated cells were seeded at 1.5 × 10^4 cells per well, with three technical replicates per group. After VEGF-A165 treatment (10 ng/mL), tube formation was evaluated after 6 h under an inverted microscope (ECLIPSE Ts2, Nikon, Japan), and the numbers of branch points and total segment lengths were quantified.

### Cell scratch assay

HBMECs were seeded into 6-well plates and grown to confluence. A linear wound was created with a sterile 200-μL pipette tip. Detached cells were removed by washing with phosphate-buffered saline (PBS), and HBMECs were then cultured in the indicated treatment medium at 37°C with 5% CO2. Images were obtained at 0 and 24 h using an inverted microscope (ECLIPSE Ts2, Nikon, Japan), and relative migration ability was quantified.

### Transwell migration assay

HBMECs were digested and resuspended in serum-free medium at a density of 2.5 × 10^5 cells/mL. A total of 100 μL of cell suspension was added to the upper chamber, whereas 600 μL of medium containing 10% FBS was added to the lower chamber. After incubation at 37°C for 24 h, the inserts were washed with phosphate-buffered saline (PBS), and migrated cells were fixed with methanol (10014118; Sinopharm Chemical Reagent Co. [SCRC], Shanghai, China) for 30 min and stained with 0.1% crystal violet staining solution (1%, G1062; Solarbio Science & Technology, Beijing, China) for 15 min. Migrated cells on the lower surface of the membrane were counted in randomly selected fields under an inverted microscope (ECLIPSE Ts2, Nikon, Japan).

### Western blot analysis

Total protein was extracted using radioimmunoprecipitation assay (RIPA) buffer (KGB5203; KeyGEN BioTECH, Nanjing, China) supplemented with phenylmethylsulfonyl fluoride (PMSF) (97064-898; Amresco, VWR International, OH, USA), and protein concentrations were measured using a bicinchoninic acid protein assay kit (KGB2101; KeyGEN BioTECH, Nanjing, China). Samples were mixed with Loading Buffer (WB2001; NCM Biotech, Suzhou, China), denatured, separated by sodium dodecyl sulfate-polyacrylamide gel electrophoresis (SDS-PAGE), and transferred to 0.2 μm polyvinylidene fluoride (PVDF) membranes (ISEQ00010; Millipore, Schwalbach, Germany). Electrophoresis and transfer were performed using a Criterion electrophoresis system and Trans-Blot transfer system (Bio-Rad, USA). Enhanced chemiluminescence (ECL) detection was carried out using ECL reagent (32209; Thermo Fisher Scientific, Pittsburgh, PA, USA), and the protein marker used was StarSignal Western Protein Marker (M227-01; GenStar, Beijing, China). Images were acquired using a Tanon 5200 imaging system (Tanon, China). The anti-SULF1 antibody (27438-1-PBS) was purchased from Proteintech (Wuhan, China); anti-vascular endothelial growth factor receptor 2 (VEGFR2) (AF6281), anti-phosphorylated VEGFR2 (p-VEGFR2) (Y1175; AF4426), anti-extracellular signal-regulated kinase 1/2 (ERK1/2) (AF0155), and anti-phosphorylated ERK1/2 (p-ERK1/2) (Thr202/Tyr204; AF1015) antibodies were purchased from Affinity Biosciences (OH, USA); anti-protein kinase B (AKT) (9272) and anti-phosphorylated AKT (p-AKT) (Ser473; 9271) antibodies were purchased from Cell Signaling Technology (Massachusetts, USA); and anti-β-actin (ab8227) together with the corresponding secondary antibodies were purchased from Abcam (Cambridge, UK). Band intensities were quantified using Image Pro Plus 6.0 software.

### qRT-PCR

Total RNA was extracted from treated HBMECs using TRIzol reagent (9109; Takara, Japan). RNA concentration and purity were measured using a NanoDrop 2000 spectrophotometer (Thermo Fisher Scientific, USA). Reverse transcription was performed using the iScript cDNA synthesis kit (1708891EDU; Bio-Rad, Hercules, CA, USA) on an Applied Biosystems Veriti 96-Well Thermal Cycler (Life Technologies, USA). qRT-PCR amplification was performed using TB Green Premix Ex Taq (Tli RNase H Plus; RR420A; Takara, Japan) on an ABI 7500 real-time PCR system (Applied Biosystems, USA). PCR primers were synthesized by Sangon Biotech (Shanghai, China). The target genes included SULF1, ANGPT2, ESM1, KDR, ICAM1, and VCAM1. Relative mRNA expression levels were calculated using the 2^-ΔΔCt method with β-actin as the internal reference.

### Statistical analysis

Statistical analyses and graph generation for cell experiments were performed using GraphPad Prism 9 (version 9.4.0; GraphPad Software, USA). Figure assembly was performed using Adobe Illustrator 2022 (version 26.3.0; Adobe, USA). Data are presented as mean ± standard deviation. Group comparisons were performed using one-way analysis of variance or two-way analysis of variance as appropriate. A two-sided P value < 0.05 was considered statistically significant.

## Results

### MMD vascular tissue exhibits intimal hyperplasia and altered spatial cellular architecture

Vascular specimens were obtained from patients with MMD (n=6), whereas control specimens were collected from patients undergoing epilepsy surgery (n=3). Multiplex immunofluorescence showed that MMD vascular tissue exhibited a thicker intimal layer and altered vessel wall morphology compared with the control group in the αSMA- and CD31-stained sections (Figure 1A–B). Quantitative analysis further showed a higher intima-to-media area ratio in MMD than in the control group (Figure 1F). We next applied Xenium in situ spatial transcriptomics to profile these vascular specimens and resolve their spatial cellular architecture. UMAP visualization identified eight major annotated cell populations in MMD and control tissues, including endothelial cells ( EC) and smooth muscle cells (SMC), and outlined the overall cellular organization represented in the profiled sections (Figure 1C–D). The dot plot showed that representative marker expression supported the annotation of these major cell populations (Figure 1E).

**Figure 1.**
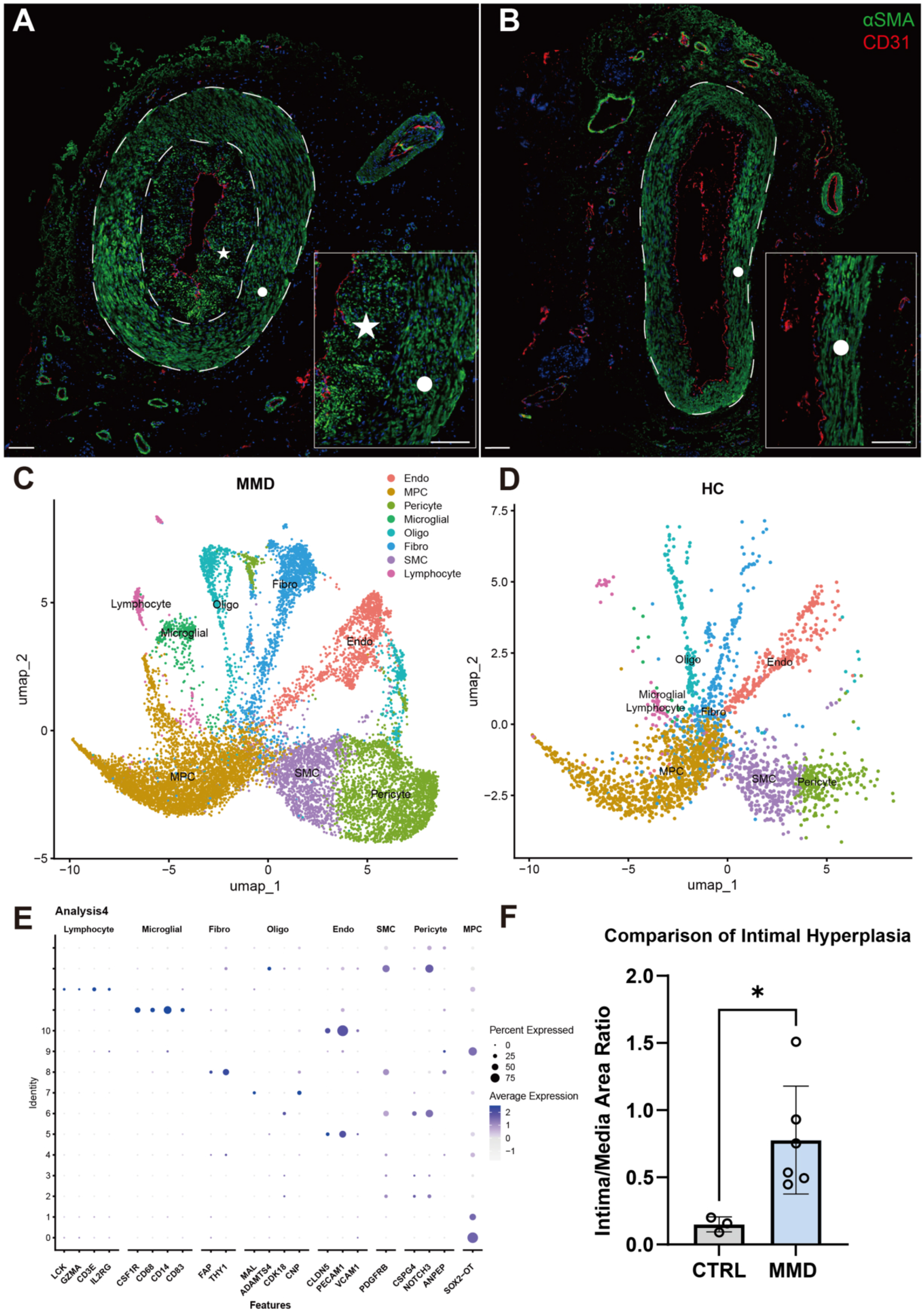
Xenium profiling revealed intimal hyperplasia and altered spatial cellular architecture in MMD vascular tissue. **A** and **B**, Representative multiplex immunofluorescence images of vascular sections from MMD and control group samples stained for αSMA and CD31. Insets show higher-magnification views of the indicated regions. Dashed lines outline the vessel wall. **C** and **D**, UMAP plots showing the major annotated cell populations in MMD and control group samples. **E**, Dot plot of representative marker genes used for cell-type annotation. Dot size indicates the percentage of cells expressing each marker, and color indicates the average expression level. **F**, Quantitative analysis of the intima/media area ratio in control group and MMD vascular samples. Results were mean ± SD. ^*^*P*<0.05.

### Differential expression and enrichment analyses revealed extracellular matrix associated transcriptional alterations

Differential expression analysis based on Xenium in situ spatial transcriptomics identified a distinct set of lesion-associated genes in MMD vascular tissue, among which SULF1 was significantly upregulated and selected for further study (log^2^FC = 2.68685, adjusted P = 1.38052E-10) (Figure 2A). Kyoto Encyclopedia of Genes and Genomes (KEGG) enrichment analysis showed that these genes were mainly enriched in pathways related to adhesion, extracellular matrix (ECM) regulation, proteoglycans, and associated signaling programs (Figure 2B). Gene Ontology (GO) enrichment analysis indicated that these transcriptional alterations were concentrated in biological processes related to cell-matrix interaction, cellular components associated with ECM organization, and molecular functions involving matrix-associated and heparin binding features (Figure 2C–E). Network visualization showed extensive gene sharing across these enriched categories, whereas clustered enrichment analysis grouped these terms into closely related functional modules (Supplementary Figures S3–S4). These results indicated that lesion-associated transcriptional alterations in MMD were concentrated in ECM-related programs.

**Figure 2.**
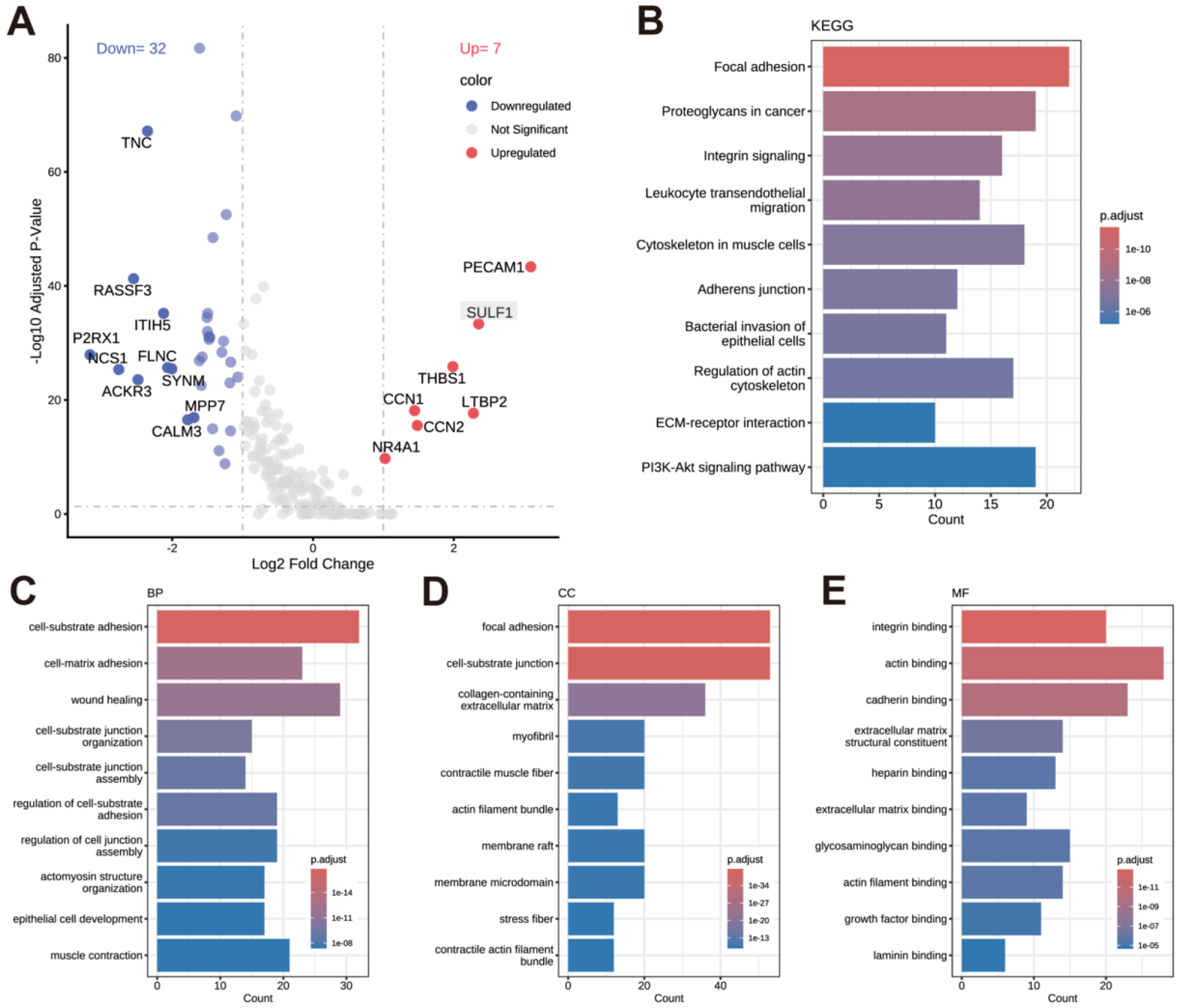
Differential expression and enrichment analyses found adhesion, extracellular matrix, and proteoglycan related programs in MMD vascular tissue. **A**, Volcano plot of differentially expressed genes between the compared groups. Representative upregulated and downregulated genes are labeled. **B**, KEGG enrichment analysis of the differentially expressed genes. **C** through **E**, Gene Ontology enrichment analyses for biological process (BP, C), cellular component (CC, D), and molecular function (MF, E). Bar length indicates gene count, and color indicates adjusted *P* value.

### Validation of SULF1 knockdown and overexpression models in HBMECs

To support the downstream signaling and functional assays, SULF1 knockdown and overexpression models were established and validated in HBMECs. Multiple knockdown constructs were first screened in HBMECs, and both Western blot and qRT-PCR demonstrated effective reduction of SULF1 expression. Among these constructs, LV-SULF1-RNAi (PSC60698-1) was selected for subsequent experiments (Figure S1). In parallel, transfection with the LV-SULF1 (KL25472-1) construct increased SULF1 expression at both the protein and mRNA levels, confirming successful overexpression in HBMECs (Figure S2). These validated HBMEC models were subsequently used for the downstream signaling and functional analyses.

### SULF1 knockdown attenuated VEGF-A165-induced activation of VEGFR2 and downstream ERK/AKT signaling

HBMECs transfected with LV-RNAi-NC or LV-SULF1-RNAi were cultured overnight in medium containing 1% FBS and then stimulated with VEGF-A165 (50 ng/mL) for 0, 10, and 30 min. Western blot analysis showed that, compared with the LV-RNAi-NC group, the LV-SULF1-RNAi group exhibited lower levels of p-VEGFR2, p-ERK1/2, p-AKT, and SULF1 across the indicated time points (Figure 3A–H). In contrast, the total protein levels of VEGFR2, ERK1/2, and AKT showed no obvious difference between the two groups. These results suggested that SULF1 knockdown may attenuate VEGF-A165-induced activation of VEGFR2 and downstream ERK/AKT signaling in HBMECs.

**Figure 3.**
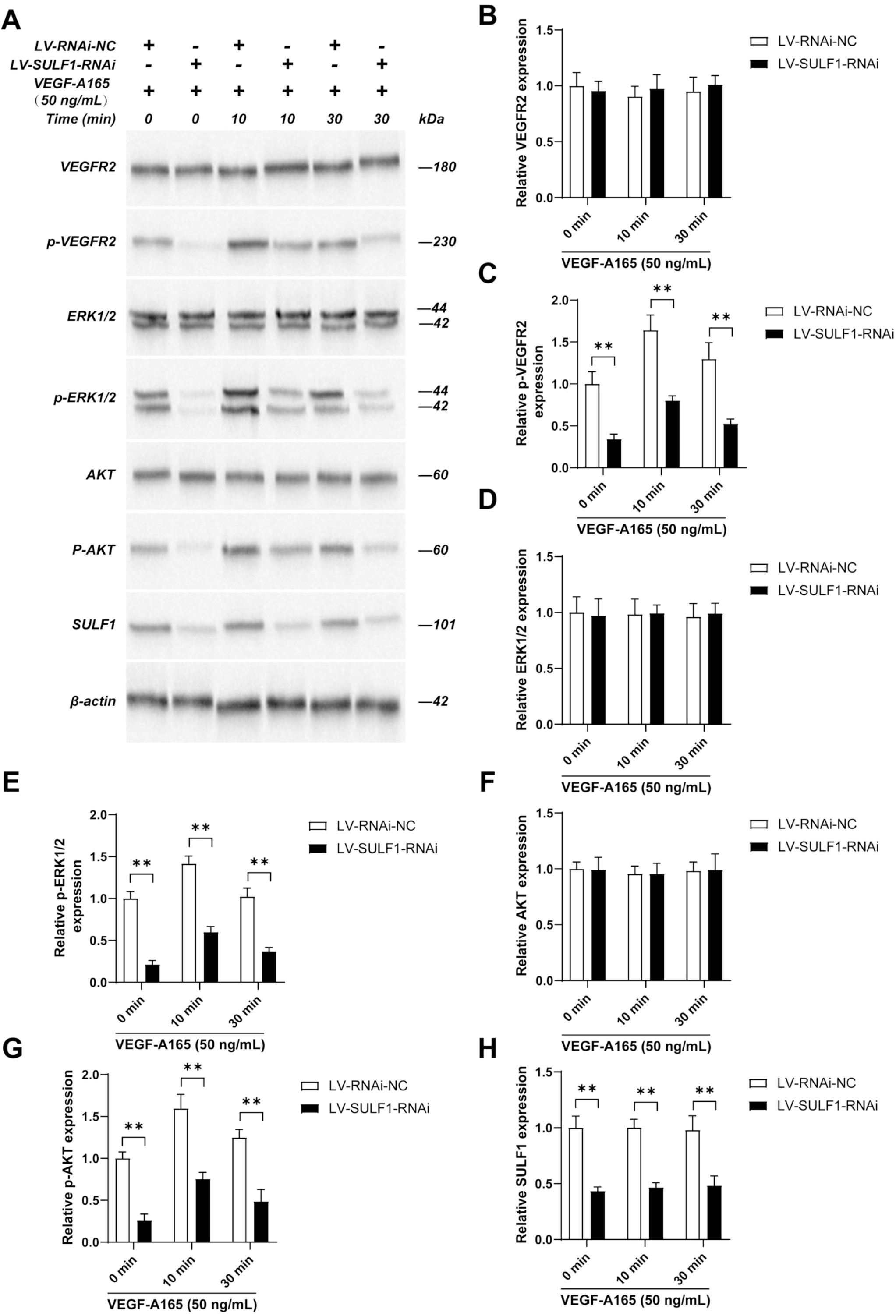
SULF1 knockdown inhibited VEGF-A165-induced activation of VEGFR2 and downstream ERK/AKT signaling in HBMECs. HBMECs were infected with LV-RNAi-NC or LV-SULF1-RNAi and cultured overnight in medium containing 1% FBS, followed by stimulation with VEGF-A165 (50 ng/mL) for 0, 10, and 30 min. **A**, Representative western blots of VEGFR2, p-VEGFR2, ERK1/2, p-ERK1/2, AKT, p-AKT, SULF1, and β-actin. **B** through **H**, Quantitative analysis of VEGFR2 (**B**), p-VEGFR2 (**C**), ERK1/2 (**D**), p-ERK1/2 (**E**), AKT (**F**), p-AKT (**G**), and SULF1 (**H**). Results were mean ± SD for three individual experiments. *n*=3, ^**^*P*<0.01.

### SULF1 overexpression was associated with enhanced VEGF-A165-induced signaling responses

HBMECs were assigned to a vector control group, a vector control group with heparinase III pretreatment, a SULF1 overexpression group, and a SULF1 overexpression group with heparinase III pretreatment. Cells were maintained in medium containing 1% FBS and then stimulated with VEGF-A165 (50 ng/mL) for 0, 10, and 30 min. Western blot analysis showed that, compared with the vector control group, the SULF1 overexpression group exhibited increased levels of p-VEGFR2, p-ERK1/2, p-AKT, and SULF1 across the indicated time points (Figure 4A–H). In contrast, the total protein levels of VEGFR2, ERK1/2, and AKT showed no obvious difference among groups. After heparinase III pretreatment, the phosphorylation responses were lower than those in the corresponding untreated groups. These results suggested that SULF1 overexpression may be associated with enhanced VEGF-A165-induced activation of VEGFR2 and downstream ERK/AKT signaling in HBMECs.

**Figure 4.**
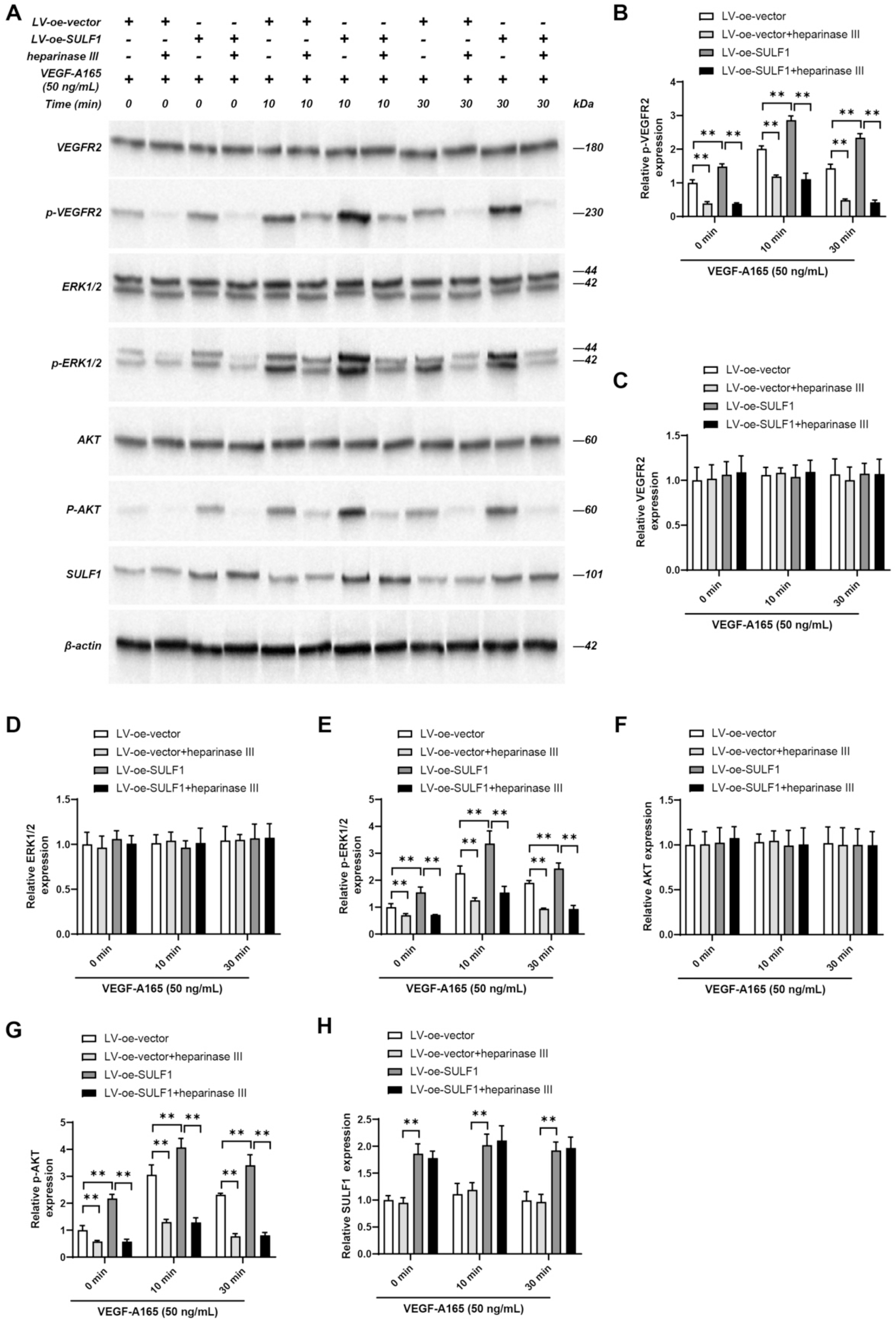
SULF1 promoted VEGF-A165-induced signaling activation in HBMECs, and heparinase III attenuated this effect. HBMECs were processed as follows: LV-oe-vector group, cells were transfected with the control lentiviral plasmid; LV-oe-vector+heparinase III group, cells were transfected with the control lentiviral plasmid and pretreated with heparinase III; LV-oe-SULF1 group, cells were transfected with the LV-SULF1 lentiviral plasmid; LV-oe-SULF1+heparinase III group, cells were transfected with the LV-SULF1 lentiviral plasmid and pretreated with heparinase III. All groups were stimulated with VEGF-A165 (50 ng/mL) for 0, 10, and 30 min. **A**, Representative western blots of VEGFR2, p-VEGFR2, ERK1/2, p-ERK1/2, AKT, p-AKT, SULF1, and β-actin. **B** through **H**, Quantitative analysis of p-VEGFR2 (**B**), VEGFR2 (**C**), ERK1/2 (**D**), p-ERK1/2 (**E**), AKT (**F**), p-AKT (**G**), and SULF1 (**H**). Results were mean ± SD for three individual experiments. *n*=3, ^**^*P*<0.01.

### SULF1 expression was associated with altered angiogenic and migratory phenotypes in HBMECs

HBMECs in the LV-RNAi-NC, LV-SULF1-RNAi, LV-oe-vector, and LV-oe-SULF1 groups were treated with VEGF-A165 (10 ng/mL) after transfection and then subjected to tube formation, transwell migration, and cell scratch assays. In the tube-formation assay, assessed after 6 h of stimulation, the LV-SULF1-RNAi group showed reduced angiogenic activity compared with the LV-RNAi-NC group, whereas the LV-oe-SULF1 group showed enhanced tube formation relative to the LV-oe-vector group (Figure 5A–C). In the transwell migration assay, assessed after 24 h, migrated cell number was reduced in the LV-SULF1-RNAi group and increased in the LV-oe-SULF1 group relative to their corresponding control groups (Figure 5D–E). In the cell scratch assay, relative migration ability showed the same directional changes, with reduced migration in the knockdown group and increased migration in the overexpression group (Figure 6A–B). These results suggested that changes in SULF1 expression were associated with altered VEGF-A165-induced angiogenic and migratory phenotypes in HBMECs.

**Figure 5.**
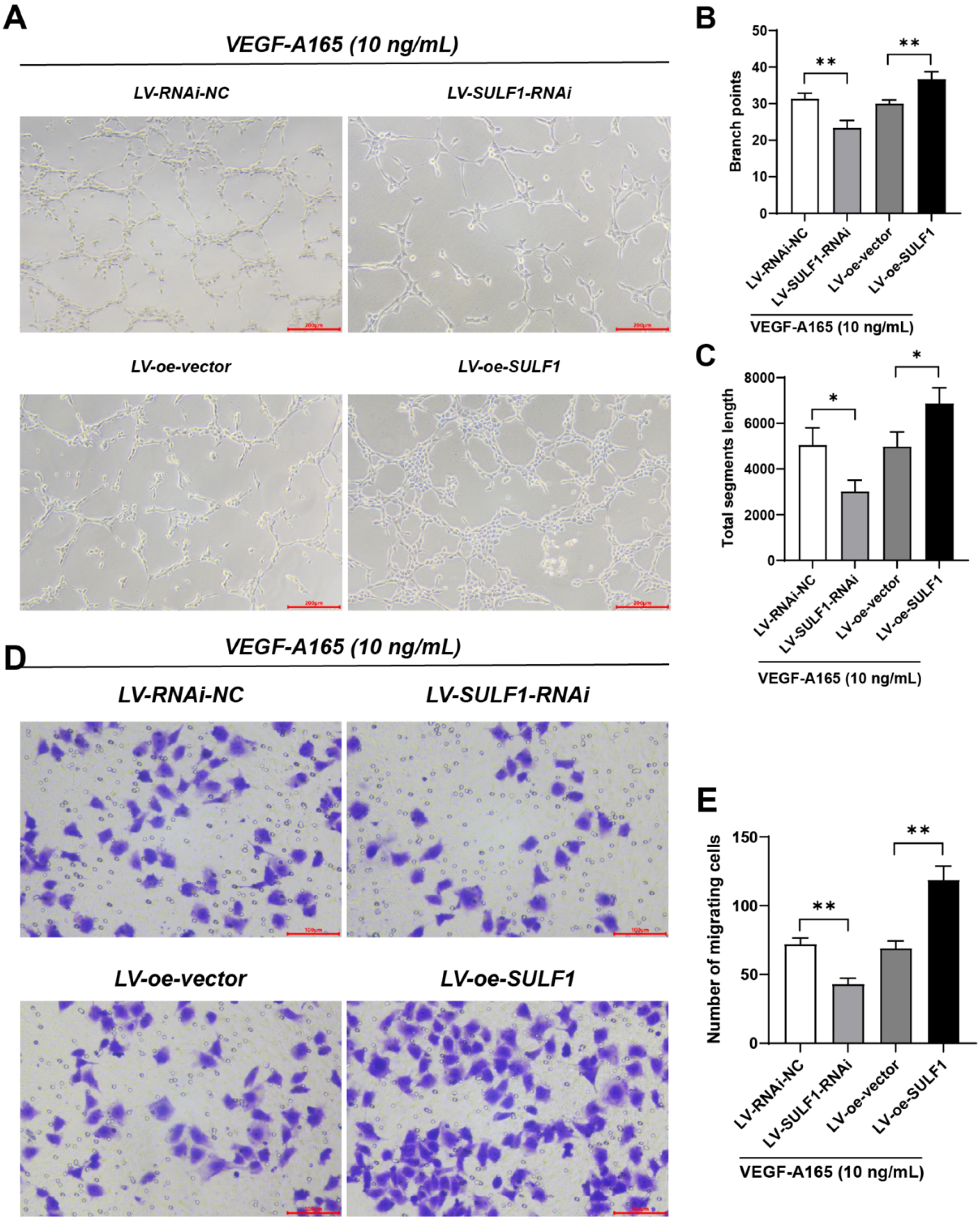
SULF1 modulated VEGF-A165-induced tube formation and Transwell migration in HBMECs. HBMECs were transfected with LV-RNAi-NC, LV-SULF1-RNAi, LV-oe-vector, and LV-oe-SULF1 lentiviral plasmids, respectively. After transfection, cells were treated with VEGF-A165 (10 ng/mL). **A**, Representative images of tube formation in each group after 6 h. Scale bars are shown in the panels. **B** and **C**, Quantitative analysis of branch points (**B**) and total segments length (**C**). **D**, Representative microscopic images of migrating HBMECs in the Transwell assay after 24 h. Scale bars are shown in the panels. **E**, Quantitative analysis of the number of migrating cells. Results were mean ± SD for three individual experiments. *n*=3,^*^*P*<0.05, ^**^*P*<0.01.

**Figure 6.**
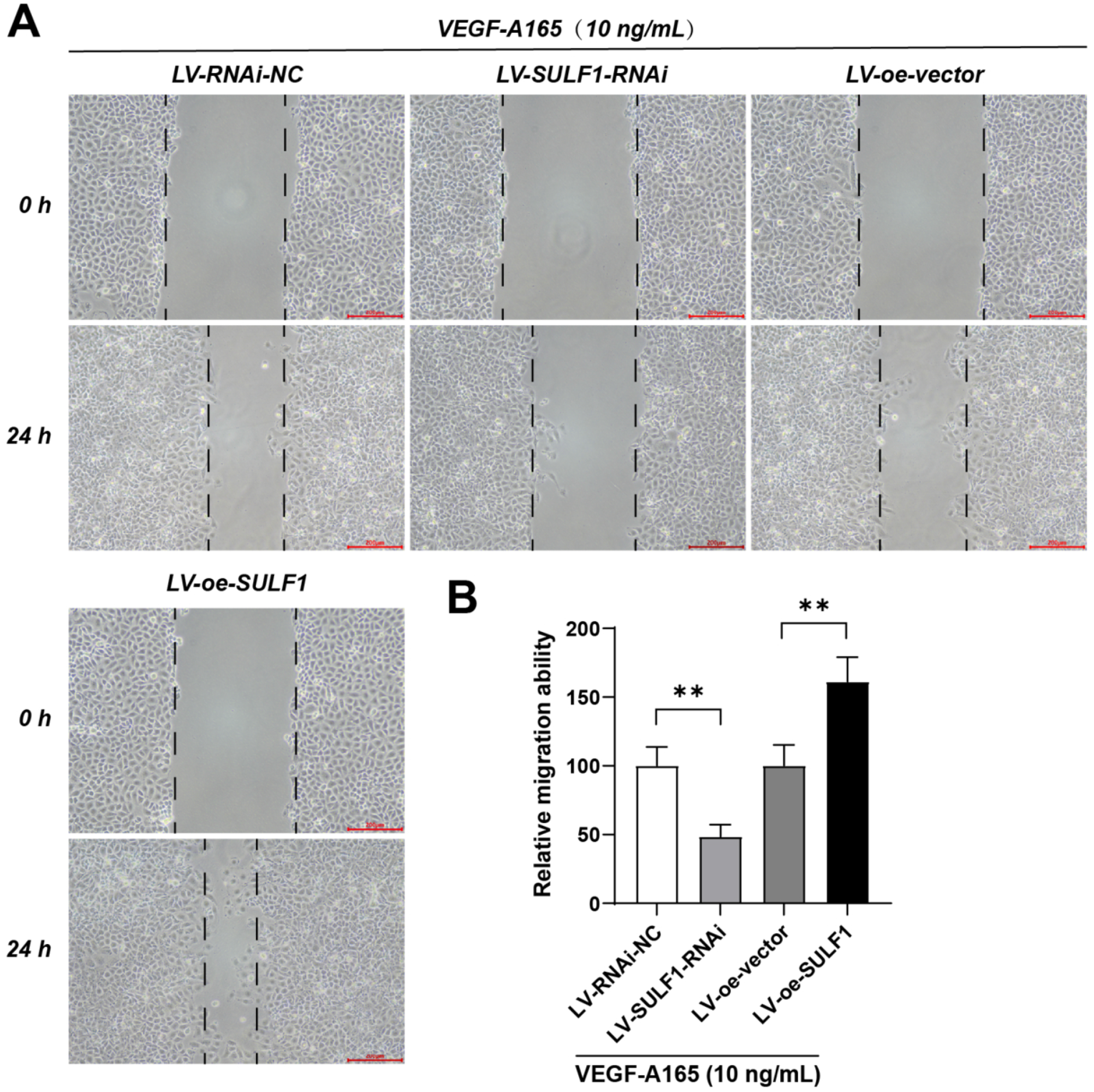
SULF1 modulated VEGF-A165-induced cell scratch migration in HBMECs. HBMECs were transfected with LV-RNAi-NC, LV-SULF1-RNAi, LV-oe-vector, and LV-oe-SULF1 lentiviral plasmids, respectively. After transfection, cells were treated with VEGF-A165 (10 ng/mL) for 24 h. **A**, Representative cell scratch images of HBMECs at 0 and 24 h in each group. Dashed lines indicate the wound margins. Scale bars are shown in the panels. **B**, Quantitative analysis of relative migration ability. Results were mean ± SD for three individual experiments. *n*=3, ^**^*P*<0.01.

### SULF1 expression was associated with transcriptional changes in angiogenesis- and adhesion-related genes

After transfection, HBMECs were assigned to the LV-RNAi-NC, LV-SULF1-RNAi, LV-oe-vector, and LV-oe-SULF1 groups and exposed to VEGF-A165 (10 ng/mL). qRT-PCR analysis was then performed after 24 h to assess transcriptional changes in angiogenesis- and adhesion-related genes. Compared with the LV-RNAi-NC group, the LV-SULF1-RNAi group showed lower mRNA levels of ANGPT2, ESM1, KDR, ICAM1, and VCAM1. In contrast, the LV-oe-SULF1 group showed higher transcript levels of these genes than the LV-oe-vector group, and the same directional pattern was observed across all five genes examined (Figure 7A–E). These results suggested that changes in SULF1 expression may be associated with altered transcriptional levels of angiogenesis- and adhesion-related genes in HBMECs under VEGF-A165 stimulation.

**Figure 7.**
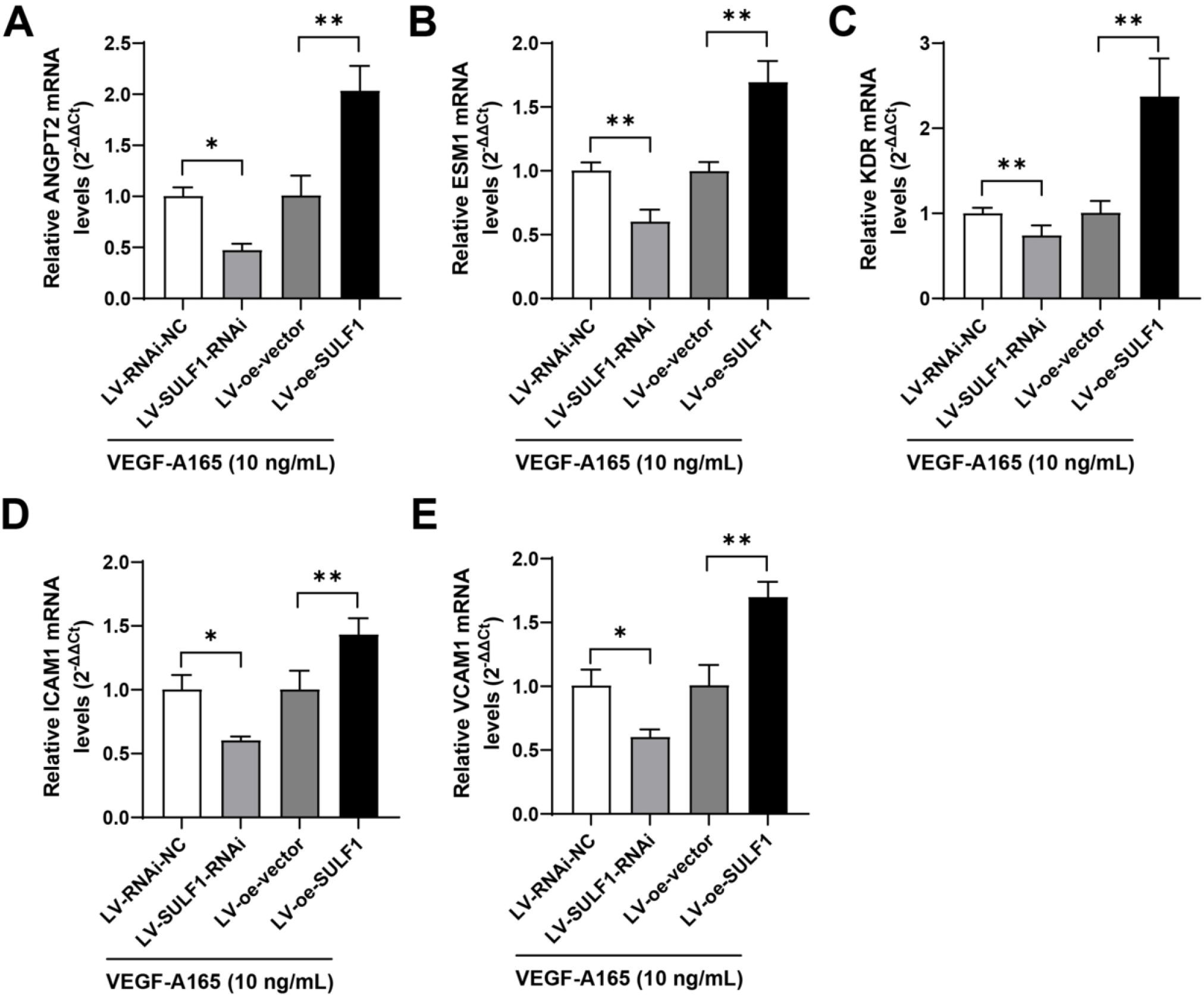
SULF1 modulated VEGF-A165-induced expression of angiogenesis- and adhesion-related genes in HBMECs. HBMECs were transfected with LV-RNAi-NC, LV-SULF1-RNAi, LV-oe-vector, and LV-oe-SULF1 lentiviral plasmids, respectively. After transfection, cells were treated with VEGF-A165 (10 ng/mL) for 24 h. The mRNA levels of ANGPT2 (**A**), ESM1 (**B**), KDR (**C**), ICAM1 (**D**), and VCAM1 (**E**) were detected by qRT-PCR. Results were mean ± SD for three individual experiments. *n*=3, ^*^*P*<0.05, ^**^*P*<0.01.

## Discussion

MMD is a chronic progressive occlusive cerebrovascular disorder characterized by progressive stenosis or occlusion of the terminal internal carotid artery and its proximal branches, together with the formation of an abnormal collateral vascular network at the base of the brain. The affected vessels typically exhibit fibrocellular intimal thickening, irregular or duplicated internal elastic lamina, and medial thinning, indicating that MMD is not merely a disease of luminal narrowing but a dynamic vasculopathy involving complex vessel-wall remodeling. Although previous genetic, epigenetic, exosomal RNA, serum proteomic, and single-cell studies have implicated vascular remodeling, dysregulated angiogenesis, and vascular-cell heterogeneity in MMD, most of these observations have remained at the level of molecular alteration or cellular composition, with limited direct information on how these abnormalities are spatially organized within the diseased vessel wall and coupled to local structural change. In the present study, by integrating histological assessment with Xenium in situ spatial transcriptomics, we found that MMD vascular tissue exhibited evident intimal hyperplasia and altered spatial cellular architecture, and that lesion-associated expression changes were mainly enriched in extracellular matrix-, adhesion-, and proteoglycan-related pathways, with prominent upregulation of SULF1. When interpreted together with the in vitro data, our findings further showed that changes in SULF1 expression were directionally associated with VEGF-A165-induced VEGFR2/ERK/AKT activation, endothelial migration, tube formation, and related gene expression changes, whereas heparinase III pretreatment attenuated these effects. Collectively, these results suggest that spatial structural disorganization, extracellular matrix–related expression changes, and SULF1-associated enhancement of endothelial VEGF responsiveness may be linked within a common lesion framework in the MMD vessel wall.

The histological and spatial findings of this study are consistent with the previous classical vascular pathological description of MMD. Previous studies have shown that affected vessels in MMD typically display fibrocellular intimal thickening, abnormalities of the internal elastic lamina, and medial thinning, indicating persistent vessel-wall remodeling rather than simple luminal narrowing alone^1,2,17^. Recent work has further advanced this field, but its main focus still falls into several directions, including histopathological features of the diseased vessel wall, endothelial-associated molecular abnormalities and their angiogenic effects, and vascular biological changes linked to genetic susceptibility. Abumiya and Fujimura^17^ pointed out that the central problem in MMD is not merely luminal narrowing, but also alteration of the intrinsic vessel-wall structure and its response to injury. Sung et al.^6^ reported that hypomethylation of the SORT1 promoter was associated with enhanced angiogenic activity, whereas Ye et al.^8^, by combining animal models, cell experiments, and transcriptomic analyses, suggested that RNF213 loss of function is related to pathological angiogenesis. In parallel, Xu et al.^18^ proposed from bulk RNA sequencing (bulk RNA-seq) of middle cerebral arteries in adult MMD that extracellular matrix-related alterations and mitochondrial dysfunction may represent major molecular features, and He et al.^11^ showed from single-cell transcriptomics of the superficial temporal artery that diseased vessels contain altered vascular-cell composition and immune-related interactions. These studies support the view that MMD vessels are not static stenotic structures, but rather a dynamic pathological environment accompanied by ongoing local cellular and molecular change. At the same time, each line of evidence has its own limitations: conventional pathology can summarize the overall morphology of vessel-wall abnormalities but cannot localize molecular alterations and their cellular context within the same tissue section; studies based on circulating samples, epigenetic readouts, or bulk RNA-seq is useful for identifying global abnormalities but cannot directly define where these changes occur within the diseased vessel wall; and dissociation-based single-cell analysis can resolve shifts in cellular composition and state but inevitably loses the original spatial neighborhood relationships. By contrast, Campos et al.^15^ have shown in atherosclerotic coronary artery that spatial transcriptomics can help resolve localized microenvironments across vessel-wall layers, and Marco Salas et al.^16^ highlighted the value of Xenium for mapping local molecular distribution at subcellular resolution. Against this background, the added value of the present study does not lie in generating another list of differential molecules, but in placing expression abnormalities back into the actual structural context of the diseased vessel wall, thereby allowing local cellular architecture, spatial localization, and molecular change to be interpreted together^19^. Against this background, the added value of the present study does not lie in generating another list of differential molecules, but in placing expression abnormalities back into the actual structural context of the diseased vessel wall, thereby allowing local cellular architecture, spatial localization, and molecular change to be interpreted together.

The Xenium data suggest that abnormalities in the diseased vessel wall are concentrated in terms related to extracellular matrix (ECM) organization, cell–matrix interaction, and proteoglycan biology, rather than being scattered across multiple unrelated directions. Differential expression analysis identified 32 downregulated and 7 upregulated genes in MMD tissue, whereas enrichment converged on focal adhesion, proteoglycans, integrin signaling, ECM-receptor interaction, leukocyte transendothelial migration, and PI3K-Akt signaling. GO terms such as heparin binding, glycosaminoglycan binding, and ECM binding pointed in a similar direction. Taken together, these findings indicate that local structural abnormalities in the diseased vessel wall are accompanied by gene-expression changes related to the ECM and proteoglycans. Xu et al.^18^ similarly regarded ECM-related alterations as an important feature in bulk RNA-seq of adult MMD middle cerebral arteries. Within this context, SULF1 deserves particular attention not only because it showed relatively high fold change and statistical significance among the upregulated genes, but also because its known biology is relevant to the enriched terms identified here. Morimoto-Tomita et al.^20^ demonstrated that SULF1 belongs to the extracellular 6-O-endosulfatases and acts on heparan sulfate chains on heparan sulfate proteoglycans (HSPGs), and Ai et al.^21^ suggested that this enzyme class can alter the sulfation state of cell-surface heparan sulfate. These molecular properties fit well with the enrichment terms involving heparin binding and glycosaminoglycan binding. The present Xenium data therefore support the interpretation that upregulation of SULF1 may be associated with ECM- and proteoglycan-related alterations in the MMD vessel wall, whereas its specific effects on endothelial behavior need to be interpreted together with the downstream cell experiments.

To further examine whether SULF1 is associated with VEGF-A165-induced endothelial signaling, we established and validated SULF1 knockdown and overexpression models in HBMECs and then performed immunoblot analysis. Compared with the LV-RNAi-NC group, the LV-SULF1-RNAi group showed lower levels of p-VEGFR2, p-ERK1/2, and p-AKT after VEGF-A165 stimulation, whereas total VEGFR2, ERK1/2, and AKT were not obviously changed. The overexpression model showed the opposite direction, with higher phosphorylation levels in the LV-oe-SULF1 group than in the LV-oe-vector group; this increase was attenuated when cells were pretreated with heparinase III. A cautious interpretation is that these findings support the involvement of heparan sulfate (HS)-related processes in the association between SULF1 change and VEGF-A165 responsiveness. Prior studies provide context for this interpretation. Simons et al.^22^ and Koch et al.^23^ summarized the role of HS and its modified states as coreceptor-level regulators of VEGFR2 signaling. Fuster and Wang^24^ emphasized that endothelial HS is an important component in the regulation of angiogenic responses. Jakobsson et al.^25^ showed that HS can potentiate VEGFR-mediated angiogenesis, and Teran and Nugent reported that heparin/HS influences the affinity of VEGF-A–receptor complexes.^26^ Taken together, our results are more consistent with the possibility that changes in SULF1 modulate the responsiveness of HBMECs to VEGF-A165 through cell-surface HS-related mechanisms, thereby affecting downstream ERK1/2 and AKT activation. However, HS 6-O-sulfation status was not directly measured in the present study, nor was binding between VEGF-A165 and heparan sulfate proteoglycans (HSPGs) or VEGFR2 biochemically assessed. The precise steps linking SULF1, HS, VEGF-A165, and VEGFR2 therefore remain to be clarified by more direct approaches.

Beyond the signaling level, changes in SULF1 expression were also reflected in endothelial phenotype and gene expression in HBMECs. Tube formation, Transwell, and cell scratch assays showed that the LV-SULF1-RNAi group had fewer branch points, shorter total segment length, fewer migrated cells, and lower relative migration than controls, whereas the LV-oe-SULF1 group showed the opposite direction. qRT-PCR further showed that mRNA levels of ANGPT2, ESM1, KDR, ICAM1, and VCAM1 shifted in the same direction. These readouts should not be viewed in isolation. Fagiani and Christofori^27^ noted that ANGPT2 is closely related to vascular destabilization and VEGF-dependent angiogenesis. Rocha et al.^28^ reported that ESM1 is not only a VEGF-related endothelial molecule but also modulates tip-cell behavior and VEGF bioavailability. The studies by Grünewald et al.^29^ and Simons et al.^22^ indicate that KDR is an integral component of endothelial VEGF responsiveness, whereas Singh et al.^30^ and Sans et al.^31^ showed that ICAM1 and VCAM1 are closely linked to endothelial activation, leukocyte adhesion, and transendothelial migration. These changes suggest that SULF1-associated effects are not limited to VEGFR2 downstream phosphorylation, but extend to endothelial phenotypes and gene-expression changes related to migration, tube formation, activation, and leukocyte adhesion. In moyamoya disease, the vessel wall is characterized not only by intimal thickening, but also by focal abnormal neovascularization and immune-related alterations. Together with the studies by He et al.^11^ and Asselman et al.^32^, our findings suggest that MMD is not simply a stenotic vasculopathy. In this context, the upregulation of SULF1 observed in the present study is better understood as being associated with local ECM remodeling and enhanced endothelial responsiveness, rather than as evidence of a single established causal driver. At present, the evidence is largely based on correlative observations and in vitro experiments, and its relevance to lesions in MMD still requires validation in disease-relevant models.

### Limitations of the Study

Several limitations should be acknowledged. First, both the MMD and control groups were small, and the control vessels were obtained from epilepsy surgery specimens rather than ideal normal intracranial vascular controls; the present findings therefore require validation in larger cohorts. Second, Xenium is a targeted spatial transcriptomic platform based on a predefined panel rather than whole-transcriptome coverage, so gene-expression changes outside the panel were not assessed in this study. Third, the functional experiments were performed mainly in the HBMEC cell line rather than in patient-derived primary endothelial cells, and the response characteristics of these models may differ from those of actual lesions. Finally, HS 6-O-sulfation status was not directly measured, and binding between VEGF-A165 and HSPGs or VEGFR2 was not directly assessed; the heparinase III results therefore support only the involvement of HS-related processes and cannot define the precise biochemical steps of SULF1 action. In addition, downstream gene changes were validated only at the mRNA level, and their protein expression and spatial distribution within MMD lesions remain to be examined.

## Conclusions

This study combined Xenium in situ spatial transcriptomics, multiplex immunofluorescence, and HBMEC-based functional assays in superficial temporal artery samples from patients with MMD, and suggests that the diseased vessel wall is characterized by intimal thickening, altered spatial cellular architecture, and gene-expression abnormalities related to the ECM, adhesion, and proteoglycans, with prominent upregulation of SULF1. The in vitro findings further suggest that changes in SULF1 expression are associated with VEGF-A165-induced signaling, migration, and tube-formation responses, whereas this association is attenuated by heparinase III treatment. These observations support the possibility that ECM-related SULF1 alterations may participate in local vessel-wall change in MMD and warrant further validation in larger cohorts and in vivo models.

## Authors’ contributions

Shihao He and Xun Ye conceived and designed the study. Shihao He, Yaoren Chang and Xiaofan Yu collected clinical samples and performed clinical data entry. Shihao He, Yaoren Chang, Xiaofan Yu, Talha Ahmed conducted spatial multi-omics analysis, bioinformatic analysis, and data visualization. Xun Ye and Yuanli Zhao revised the manuscript critically for important intellectual content. All authors reviewed and approved the final manuscript.

## Acknowledgement

We thank the healthcare staff in the department for their assistance with this project. We also sincerely thank all participants for their support and cooperation.

## Consent for publication

Not applicable.

## Data availability statement

The data presented in the current study are available from the corresponding author upon reasonable request.

